# Identification of Genes Required for Spatial Control and Mechanical Resilience of Cytokinesis during *Caenorhabditis elegans* Embryogenesis

**DOI:** 10.1101/2025.03.01.641011

**Authors:** Chelsey Lynn Gough, Yuxuan Rain Xiong, Aoi Hiroyasu, Taile Li, Christina Rou Hsu, Viktorija Juciute, MinJee Kim, Kalen Dofher, Kenji Sugioka

## Abstract

Cytokinesis is the final step of cell division, in which the dividing cell is physically separated into two daughter cells by the contractile ring. The contractile ring is a highly resilient molecular machine that can function properly under mechanical stress. Additionally, its function, position, and orientation are spatially modulated in developing animals to regulate morphogenesis. Although essential regulators of cytokinesis have been identified through previous genetic screens, the molecular mechanisms underlying these spatial controls and the mechanical resilience of cytokinesis remain elusive. To identify cytokinesis regulators involved in these processes, we performed a high-throughput RNAi screen using a gain-of-function mutant of actin that exhibits ectopic cortical contraction and abnormal spatial control of cytokinesis in *Caenorhabditis elegans* embryos. We obtained a list of early embryonic genes that suppress embryonic lethality in an *act-2* mutant background. Two parallel secondary screens of candidate genes were conducted. The first secondary screen in a wild-type background identified 69 candidate genes regulating spatial cytokinesis control—asymmetric ring closure, positioning, and rotation—during early embryogenesis. The second secondary screen in the *act-2(or295)* background identified four genes required for cytokinesis in this background, including microtubule regulators, *evl-20/ARL2*, and *lpin-1/Lipin1*. This study will serve as a useful resource for the development of future hypotheses and provide insights into the precise regulation of cytokinesis in tissues.

## Introduction

Cytokinesis is the final step of cell division, in which the contractile ring physically separates the dividing cell into two daughter cells (Pollard and O’Shaughnessy 2019). The contractile ring is a mechanically resilient molecular machine capable of withstanding mechanical stress and repairing physical damage (Silva *et al*. 2016; Singh *et al*. 2019). Additionally, its orientation and position are spatially regulated in three distinct ways during morphogenesis (Sugioka 2021 Dec 23). First, asymmetric contractile ring closure, often found in zygotes and epithelial tissue, is critical for maintaining cytokinesis resilience, epithelial integrity, and polarity (Maddox *et al*. 2007; Herszterg *et al*. 2013; Taulet *et al*. 2017). Second, cell rotation during cytokinesis is required for dorsal-ventral axis establishment in *C. elegans* (Priess and Thomson 1987; Schierenberg and Junkersdorf 1992) and is implicated in the establishment of animal left-right asymmetry in pond snails and *C. elegans* (Meshcheryakov and Beloussov 1975; Wood 1991; Kuroda *et al*. 2009; Naganathan *et al*. 2014). Finally, asymmetric positioning of the contractile ring during asymmetric cell division is critical for generating unequal daughter cell sizes, which is required for subsequent cell fate decisions (Jankele *et al*. 2021). Despite their significance, the regulators of mechanical resilience and the spatial control of cytokinesis remain largely unknown.

Decades of studies have revealed the essential components of the contractile ring, which are evolutionarily conserved in eukaryotes (Pollard 2017). First, the small GTPase RhoA is activated by the centralspindlin complex, thereby initiating contractile ring assembly via stimulation of downstream effectors such as formin and Rho-associated kinase (Basant and Glotzer 2018). Contractile ring function also requires structural components such as anillin (Piekny and Maddox 2010). As the contractile ring constricts through actin-myosin sliding, membrane trafficking pathways contribute to cytokinesis by delivering additional membrane to the ingressing furrow to accommodate the growing cell surface area (Frémont and Echard 2018). Finally, the daughter cells separate via ESCRT-III-mediated abscission at the midbody (Mierzwa and Gerlich 2014). Previous forward genetic and RNAi-based screens in yeast and various animal systems were typically performed in wild-type backgrounds and focused on identifying essential regulators of cytokinesis (Eggert et al. 2004; Echard et al. 2004; Sönnichsen et al. 2005). A recent study in *C. elegans* performed a small-scale candidate screen in zygotes under uniaxial compression and identified several regulators required for the mechanical resilience of cytokinesis (Singh *et al*. 2019). However, a large-scale genetic screen focusing on the physical and developmental control of cytokinesis is still needed.

In this study, we performed an RNAi screen targeting 4,218 early embryonic genes in an actin gain-of-function mutant of *C. elegans*. The *act-2(or295)* mutation lies near the ATP-binding pocket, leading to filament stabilization and hyperactive cortical contractility (Willis *et al*. 2006). We found that the spatial control of cytokinesis, such as asymmetric ring closure in the zygote, rotational cytokinesis in the 2-cell stage, and asymmetric ring positioning in the 6-cell stage, is also abnormal in this mutant. Therefore, we reasoned that RNAi knockdown that either suppresses or enhances *act-2(or295)* embryonic lethality may reveal regulators of both the spatial control and mechanical resilience of cytokinesis. Among the 689 genes that suppress *act-2(or295)* embryonic lethality, we performed two secondary screens on 385 selected candidates: (i) a screen in a wild-type background for regulators of spatial cytokinesis control; and (ii) a screen in the *act-2(or295)* background to uncover genes essential for furrow completion under heightened cortical contractility. We identified 69 candidate regulators of the spatial control of cytokinesis and 4 genes required for cytokinesis completion in an *act-2(or295)* background.

## Results

### *act-2(or295)* exhibits defects in three distinct modes of spatial control of cytokinesis

The *C. elegans* embryo is a suitable model system for studying the mechanisms underlying the spatial control of cytokinesis, as it exhibits an invariant cell lineage and three distinct modes of spatially controlled cytokinesis within an hour after fertilization (Figure 1A). In zygotes, the cleavage furrow forms asymmetrically, leading to asymmetric contractile ring closure (Figure 1A, left; Figure 1B, wild-type)(Maddox *et al*. 2007). At the 2-cell stage, the somatic blastomere AB undergoes rotation during cytokinesis (Figure 1A, middle; Figure 1C, wild-type) (Schierenberg and Junkersdorf 1992). This rotation positions the posterior daughter cell ABp on the dorsal side and is critical for the establishment of the dorsal-ventral axis (Priess and Thomson 1987; Schierenberg and Junkersdorf 1992). At the 6-cell stage, the endomesodermal precursor EMS undergoes asymmetric cell division (Goldstein 1992). We found that the contractile ring is initially formed in the equatorial region but becomes displaced posteriorly during cytokinesis (Figure 1A, right; Figure 1D, wild-type).

**Figure 1.**
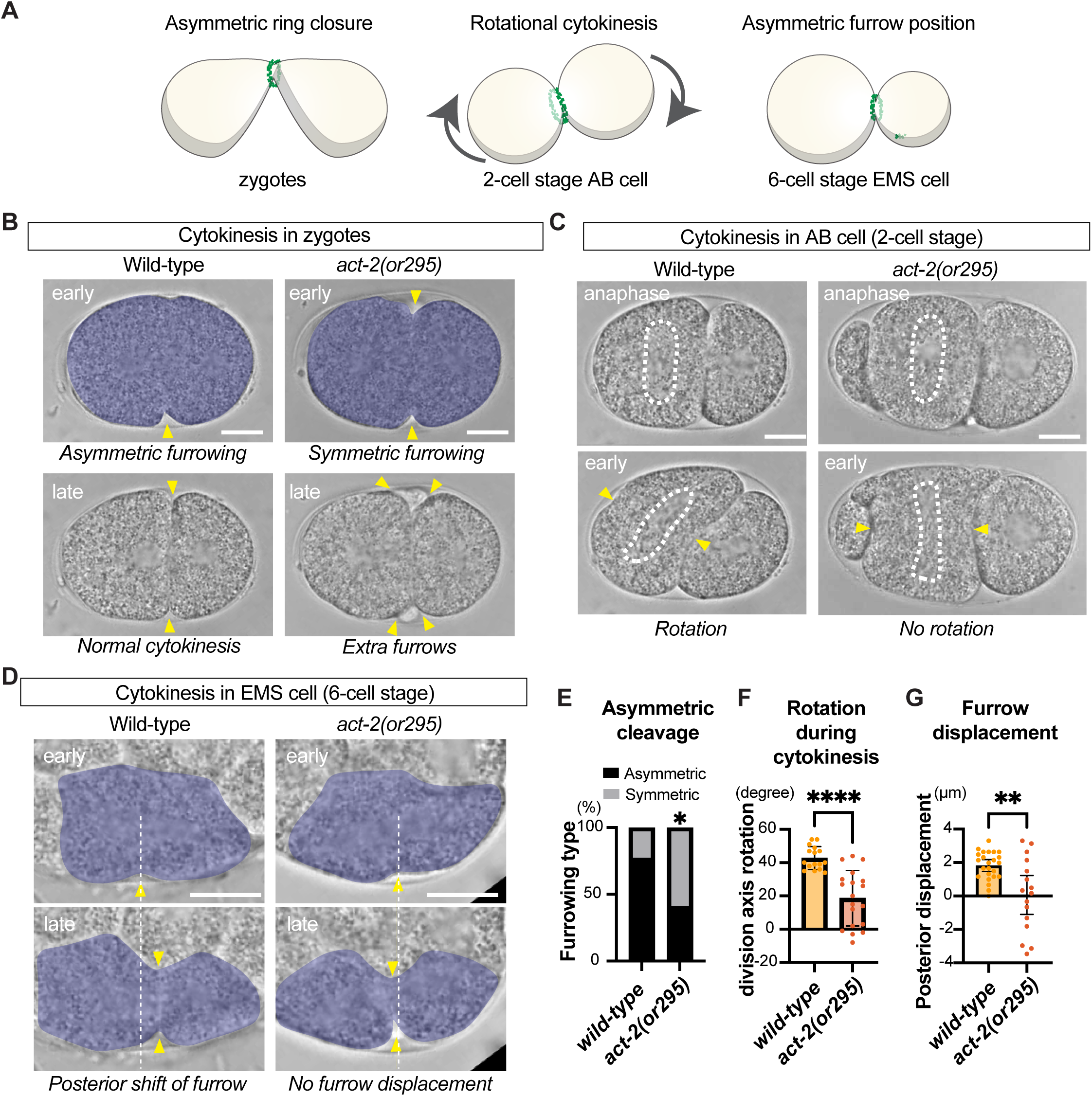
*act-2(or295)* mutant exhibits abnormal spatial control of cytokinesis. (**A**) Three modes of spatial control of cytokinesis during animal development. (**B**) Asymmetric contractile ring closure in zygotes. The contractile ring closes asymmetrically in wild-type embryos but is defective in *act-2(or295)*. *act-2(or295)* mutants also exhibit extra furrow phenotypes at later stage of cytokinesis. (**C**) Rotational cytokinesis in the AB cell. At the two-cell stage, the AB cell normally rotates during cytokinesis, but this rotation is defective in *act-2(or295)*. Dotted white lines indicate mitotic spindle position. (**D**) Asymmetric positioning of the contractile ring in the EMS cell. At the 6-cell stage, the EMS forms a cleavage furrow in the equatorial region, which then shifts towards the posterior. In *act-2(or295),* this posterior furrow shift is often defective. (**E**-**G**) Quantification of zygote furrow asymmetry (E), cellular rotation during AB cell division (F), and posterior furrow displacement during EMS cell division (G). Yellow arrowheads indicate the furrow position. Scale bars represent 10 µm. Error bars represent 95% confidence intervals. Fisher’s exact test was performed in panel E and Welch’s t-test was used for panels F and G. P-values: “****”, “**”, “*”and “ns” indicate p < 0.0001, p < 0.01, p < 0.05, and p > 0.05, respectively.

We used a temperature-sensitive and semi-dominant actin gain-of-function mutant, *act-2(or295),* which exhibits hypercontractility and stabilized actin filaments (Willis *et al*. 2006), to investigate the role of actin dynamics in the spatial control of cytokinesis. In zygotes, *act-2(or295)* exhibited symmetric furrowing more often than the wild-type (Figure 1B and 1E). Additionally, cytokinesis often resulted in extra furrow formation (Figure 1B, bottom right), suggesting that the normal cytokinesis process is also compromised by this mutation. During AB cell division at the 2-cell stage, *act-2(or295)* embryos exhibited a reduced degree of rotation during cytokinesis (Figure 1C and 1F). During EMS cell division at the 6-cell stage, *act-2(or295)* displayed an abnormal shift of the contractile ring; some embryos showed no shift in position, whereas others showed an anterior shift (Figure 1D and 1G). In summary, *act-2(or295)* mutants display defects in all three modes of spatially controlled cytokinesis and also exhibit defects in proper contractile ring formation.

### High-throughput RNAi screen in *act-2(or295)* mutant background

As shown in a previous study, *act-2(or295)* mutants exhibit ectopic cortical furrowing during cytokinesis (Figure 2A) (Willis *et al*. 2006). This ectopic furrowing was suppressed by knockdown of the actomyosin regulators such as myosin regulatory light chain (*mlc-4*), profilin (*pfn-1*), or formin (*cyk-1*) (Willis *et al*. 2006). We also observed reduced ectopic furrowing after partial knockdown of RhoA activator *ect-2(RNAi)/RhoGEF* (Figure 2A; bottom). We hypothesize that the aberrant spatial cytokinesis dynamics observed in Figure 1 and embryonic lethality in *act-2(or295)* mutants result from excessive cortical contractility, and that RNAi knockdowns that either suppress or enhance *act-2*(or295) embryonic lethality may include novel regulators of cytokinesis. According to single-cell RNA-seq data, 5,928 genes are expressed in early *C. elegans* embryos (Tintori et al., 2016). Of these, 4,445 genes are included in the Ahringer feeding RNAi library (Kamath and Ahringer, 2003), and we performed a feeding RNAi screen targeting them (Figure 2A). Both wild-type and *act-2(or295)* mutants expressing an intestinal GFP reporter (*vha-6*p::GFP) were fed dsRNA-expressing bacteria at 20°C, a temperature at which *act-2(or295)* exhibits 80.2% embryonic lethality (n > 100). Instead of performing standard feeding RNAi on Nematode Growth Media (NGM) plates, we incubated the L1-stage larvae in liquid NGM supplemented with dsRNA-expressing bacteria (hereafter “liquid-based RNAi”) in 96-well plates (Figure 2A). We then counted the number of hatched worms in each well by imaging the intestinal GFP signal using high-content screening microscopy (Figure 2B-C). The effects of RNAi on embryonic viability in both wild-type and *act-2(or295)* strains were measured by computing Z-scores using the mean and standard deviation of worm counts within each 96-well plate, rather than using the global mean and standard deviation of the entire screen, to mitigate technical variation. For each of the 4,218 analyzed genes, we obtained three parameters: (1) Z-score in the control background, (2) Z-score in the *act-2(or295)* background, and (3) the difference between the two (Figure 3A).

**Figure 2.**
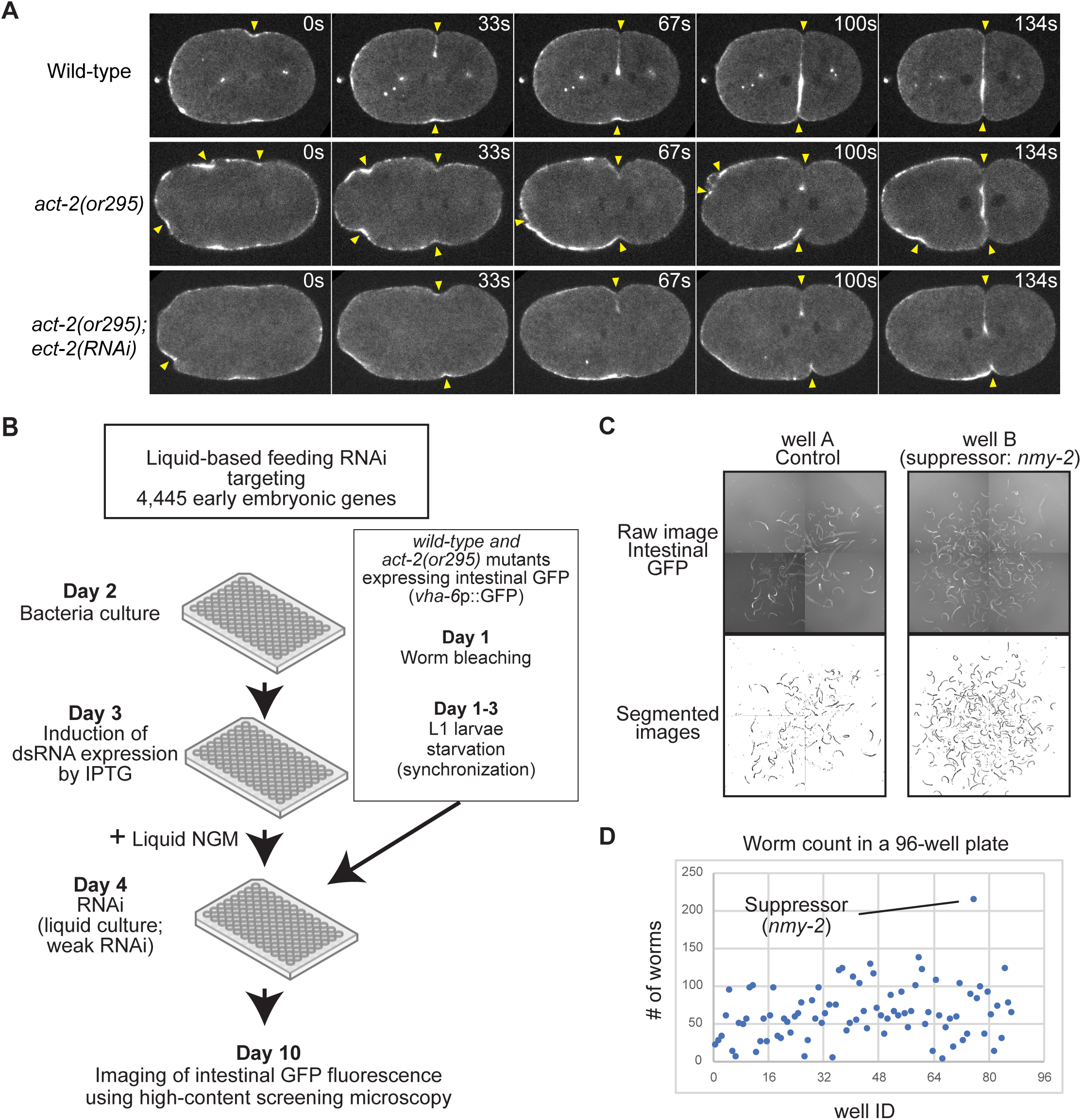
High-throughput RNAi screen using *act-2(or295)* mutants. (**A**) Ectopic cortical furrowing during cytokinesis in act-2(or295) mutants. Non-muscle myosin II::GFP localization after cytokinesis onset (0 sec). Arrowheads indicate furrows. Note that the wild-type strain also carries the centriole marker SAS-7::GFP. (**B**) Design of the RNAi screen. Wild-type and *act-2(or295)* mutants expressing an intestinal GFP reporter were bleached, and purified embryos were hatched without food to induce starvation-induced L1 larval arrest. The synchronized L1 larvae were cultured in liquid NGM containing dsRNA expressing feeding RNAi bacteria. Hatched F1 larvae were then imaged using high-content screening microscopy. (**C**) Intestinal GFP reporter signals for the control and one suppressor, *nmy-2(RNAi)*. Segmented images were used to count the number of hatched worms. (**D**) The number of hatched worms in a 96-well plate. *nmy-2(RNAi)* resulted in a significantly higher number of hatched worms, identified it as a suppressor of *act-2(or295)*.

**Figure 3.**
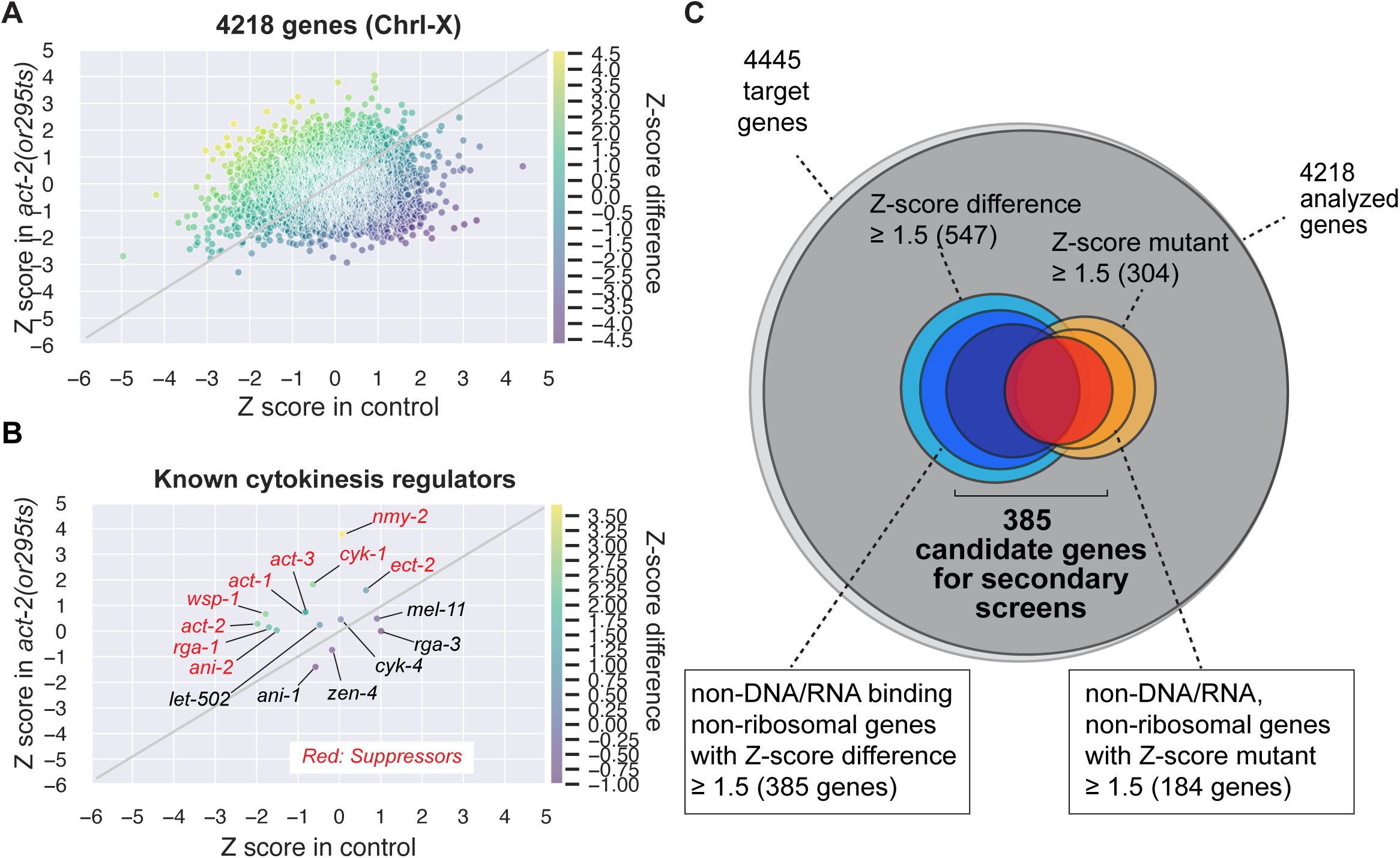
Summary of the primary RNAi screen. (**A**) Scatter plot of Z-scores for 4,218 analyzed genes. Z-scores in the control strain and *act-2* mutants are shown on the x- and y-axes, respectively. The difference in Z-scores between control and *act-2* is indicated by the color code. (**B**) Scatter plot of Z-scores highlighting known regulators of cytokinesis. As indicated by red, many of them were identified as suppressors. (**C**) Summary of identified actin suppressors. Among the non-DNA/RNA-binding and non-ribosomal genes, 385 and 184 suppressors were identified based on the Z-score difference and the Z-score in *act-2* mutants, respectively. Of the 454 suppressor genes, worm-specific genes lacking known functions or conserved domains were removed, leaving 385 genes for the secondary screens.

We found that liquid-based feeding RNAi was consistently less effective than standard plate-based feeding RNAi, possibly due to the lower density of bacteria or differences in worm feeding behavior. Because of this weaker knockdown effect, we identified the essential cytokinesis regulator *nmy-2* (non-muscle myosin II) as a suppressor of *act-2(or295)* lethality in our screen (Figure 2C). Furthermore, we identified components of the RhoA signaling pathway (*ect-2/RhoGEF*, *rga-1/RhoGAP, cyk-1/formin*), structural components of the contractile ring (*act-1/-2/-3*, *ani-2/anillin*), and a regulator of branched actin assembly (*wsp-1/N-WASP*) as suppressors (Figure 3A-B).

In this primary screen, we set a less conservative threshold and designated any RNAi with a mutant Z-score ≥ 1.5 or a Z-score difference ≥ 1.5 as a suppressor of *act-2* lethality (Figure 3C). Conversely, RNAi knockdown resulting in failed hatching or significantly reduced hatching in the *act-2(or295)* background was classified as an enhancer. Hereafter, we focus only on *act-2(or295)* suppressors, as enhancers may include genes whose knockdown additively increased the lethality of *act-2(or295)* without indicating a specific genetic interaction. We found that 304 genes displayed Z-scores ≥ 1.5 in *act-2(or295)*, and 547 genes exhibited Z-score differences ≥ 1.5 (Figure 3C; a total of 689 genes). In pursuit of genes directly regulating cytoskeletal dynamics or cell signaling, we excluded those encoding DNA-binding, RNA-binding, and ribosomal proteins (which likely affect general viability through transcription or translation). After this filtering, 385 genes had Z-score differences ≥ 1.5, and 184 genes showed Z-scores ≥ 1.5 in *act-2(or295)*, leading to a total of 454 *act-2* suppressor genes (Figure 3C). From this list, we selected genes that have a known function, a conserved domain, or are conserved in humans, yielding a final set of 385 candidate genes used for the secondary screens (Figure 3C).

### Identification of regulators of spatial control of cytokinesis

Next, we performed a secondary screen to identify regulators of the spatial control of cytokinesis among 385 genes characterized as *act-2(or295)* suppressors (Figure 4A). For each RNAi knockdown, we collected three live-imaging movies capturing embryogenesis from the 1-cell to the 7-cell stage.

**Figure 4.**
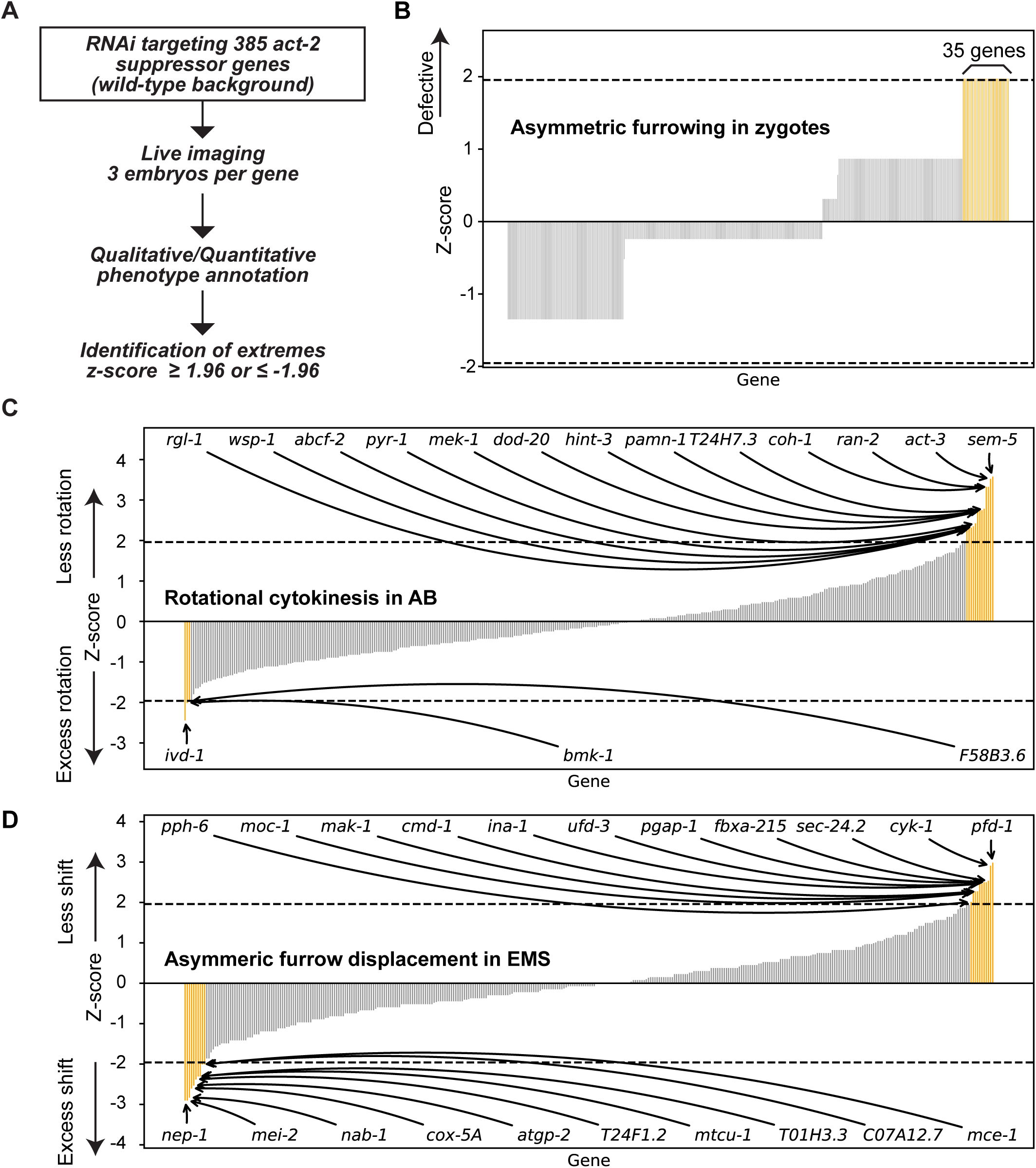
Secondary screen for spatial cytokinesis regulators. (**A**) Design of the secondary screen. A total of 385 5 *act-2* suppressor genes were targeted by standard plate-based feeding RNAi in a wild-type background. For asymmetric furrowing in zygotes, we performed a qualitative assessment, whereas for AB cell rotation and EMS cell furrow shift, we measured the degree of rotation and the extent of furrow shift, respectively. (**B**-**D**) Candidate regulators of asymmetric contractile ring closure (B), rotational cytokinesis (C), and asymmetric furrow shift (D). Orange bars indicate extreme samples with Z-scores ≥ 1.96 or ≤ -1.96.

For asymmetric contractile ring closure in zygote, we visually classified an embryo as defective if the cleavage furrow ingressed asymmetrically. Because asymmetric ring closure can occur in any orientation, furrowing asymmetry is not detectable in our 2D imaging setup when the ring closure occurs along the imaging axis. Indeed, about 23% of wild-type zygotes exhibited symmetric furrowing patterns (n = 22, Figure 1E). Thus, we aimed to identify extreme cases that displayed a high penetrance of defects. We scored phenotypes for each collected movie (score = 0 if defective, score = 1 if normal) and calculated a Z-score for each gene using the mean and standard deviation across all *act-2* suppressors. We identified 35 genes exhibiting symmetric contractile ring closure in all three movies (Z-score ≥ 1.96; Figure 4B). These candidates include *dhhc-9*/ZDHHC16, which encodes S-palmitoyltransferase, and *cyn-6*/PPIB, which encodes peptidylprolyl isomerase B (Figure 5A).

**Figure 5.**
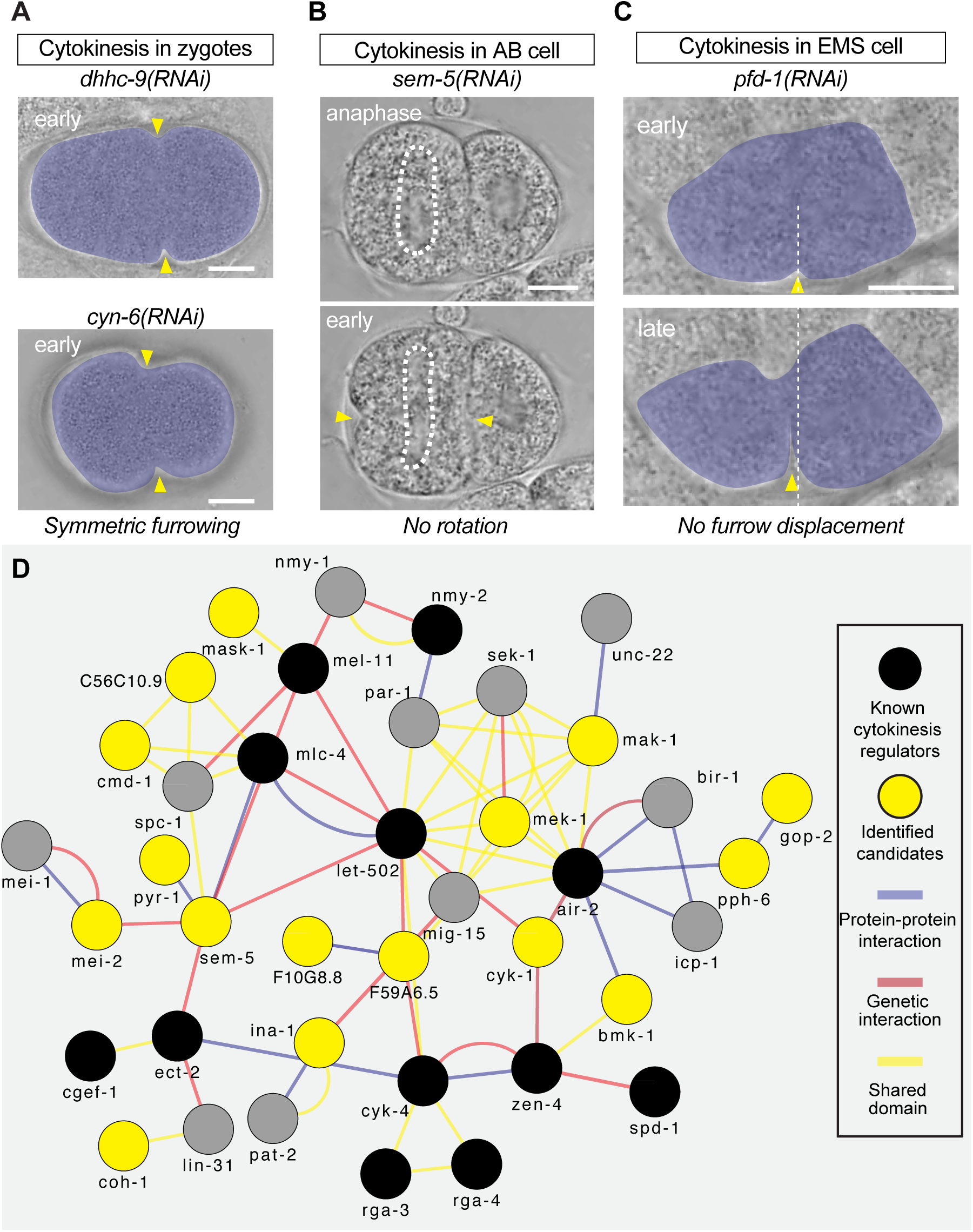
Identification of spatial cytokinesis regulators. (**A**) Defects in asymmetric furrowing in *dhhc-9(RNAi)*. (**B**) Defects in rotational cytokinesis in *sem-5(RNAi)*. (**C**) Defects in asymmetric furrow displacement in *pfd-1(RNAi)*. (**D**) Gene interaction network among identified candidate genes and known cytokinesis regulators. Protein-protein interactions (PPI), genetic interactions, shared domain information are shown. Candidates without known interactions are not displayed. Scale bar indicates 10 µm.

For the rotational cytokinesis during AB cell division, we measured the angle of AB division axis rotation. We identified 13 genes with Z-scores > 1.96 (indicating reduced rotation) and 3 genes with Z-score ≤ -1.96 (indicating excess rotation) (Figure 4C). We found that *sem-5*/Grb2, involved in epidermal growth factor signaling, causes strong defects in AB cell rotation (Figure 5B).

For the asymmetric contractile ring displacement during the EMS cell cytokinesis, we calculated the shift of the cleavage furrow along the anterior-posterior axis before and after cytokinesis for each movie. We identified 11 genes with Z-score ≥ 1.96 (indicating less furrow displacement) and 10 genes with Z-score ≤ -1.96 (indicating excessive displacement). These candidates include *pfd-1*/PFDN1, which encodes a prefoldin subunit known to be involved in actin folding (Figure 5C).

Finally, we performed a gene network analysis using the 69 identified genes and found that 16 candidate genes—either genetically or physically, and directly or indirectly— interact with known cytokinesis regulators (Figure 5D).

### Identification of regulators of mechanical resilience of cytokinesis

In *act-2(or295)* embryos, which exhibit hypercontractility, cells are under constant mechanical stress yet display only subtle defects in contractile ring formation (e.g., extra furrow shown in Figure 1B) and show no defects in cytokinesis completion in zygotes (Figure 6B). Thus, we reasoned that the genes required for cytokinesis in *act-2(or295)* may include regulators of the mechanical resilience of cytokinesis. We selected 139 genes from the 385 *act-2* suppressors based on the following criteria: for genes on Chromosome I and II, RNAi knockdown caused > 67% defects in asymmetric furrowing in the wild-type background; for genes on the remaining chromosomes, RNAi knockdown caused 100% embryonic lethality in *act-2(or295)* (Figure 6A). We then performed an RNAi screen targeting these 139 genes in an *act-2* mutant background using a standard plate-based feeding RNAi (Figure 6A). Of the 139 candidates examined, RNAi knockdown of 12 genes led to mild to severe cytokinesis defects. Among these, five genes are known cytokinesis regulators of cytokinesis: the actin genes (*act-1, act-2, act-3*), *ect-2/RhoGEF* (which activates RhoA), and *nmy-2* (non-muscle myosin II). Another three genes (*egg-6, cks-1, ubc-12*) are involved in eggshell formation, cell cycle regulation, and ubiquitination, respectively, and are known to cause cytokinesis defects when disrupted in wild-type background. Two additional genes (*tba-1*/α-tubulin and the centrosomal/centriolar component *spd-2*) function in microtubule assembly and organization.

**Figure 6.**
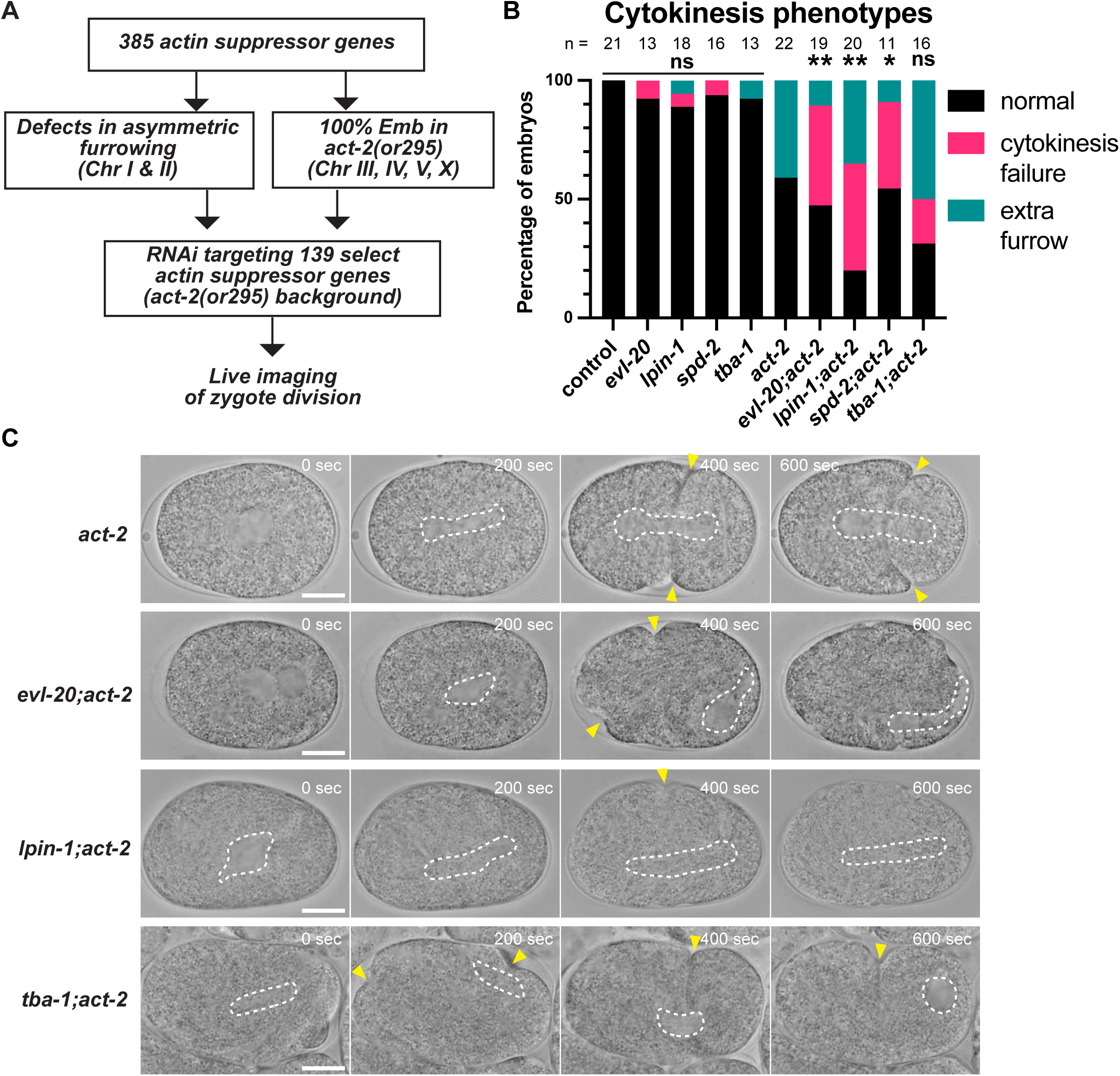
Identification of genes required for cytokinesis in the *act-2(or295)* background. (**A**) Design of the secondary screen. A total of 139 genes were selected based on defects in asymmetric furrowing in zygotes or 100% embryonic lethality in the *act-2* mutant background from 385 5 *act-2* suppressor genes. These genes were knocked-down in *act-2(or295)* mutants and tested for their requirement in cytokinesis completion. (**B**) Genes identified as required for cytokinesis completion in *act-2(or295)* mutants. P-values were calculated by chi-square test, followed by Holm-Sidak adjustment for multiple comparisons. P-values: “**”, “*”, and “ns” indicate p < 0.01, p < 0.05, and p > 0.05, respectively. (**C**) Cytokinesis phenotype after candidate gene knockdown. White dotted lines indicate the mitotic spindle position, and yellow arrowheads indicate the cleavage furrow. Scale bars, 10 µm.

The final two genes are relatively less understood. First, *evl-20/ARL2* encodes ADP-ribosylation factor-like GTPase that localizes to the periphery of astral microtubules and the plasma membrane (Antoshechkin and Han 2002). A previous study reported that *evl-20* knockdown or mutation reduces microtubule levels during embryogenesis and produces multinucleated cells later in development, such as in vulval precursor cells and the somatic gonad (Antoshechkin and Han 2002). However, the early embryonic phenotype was not described, and cytokinesis has not been analyzed in detail. *lpin-1/Lipin 1* encodes phosphatidic acid phosphatase involved in polyunsaturated fatty acids (PUFAs) production and longevity (Jung *et al*. 2020). Previous studies showed that *lpin-1* is required for proper endoplasmic reticulum organization and timely nuclear envelope breakdown during embryogenesis (Golden *et al*. 2009; Gorjánácz and Mattaj 2009). In *Toxoplasma gondii*, the loss of Lipin results in multinucleation, possibly due to cytokinesis failure (Dass *et al*. 2021).

To identify genes specifically required for cytokinesis in *act-2(or295)*, we compared RNAi phenotypes between wild-type and *act-2* backgrounds and found that *evl-20*, *lpin-1*, *spd-2*, and *tba-1* induced more severe cytokinesis defects in *act-2(or295)* (Figure 6B). The phenotypes of *spd-2(RNAi);act-2(or295)* and *tba-1(RNAi); act-2(or295)* were relatively mild, as these embryos still formed a furrow that sometimes regressed later (Figure 6C, bottom). By contrast, *evl-20(RNAi);act-2(or295)* and *lpin-1(RNAi); act-2(or295)* failed to form a fully developed furrow, suggesting that these genes are involved in contractile ring formation (Figure 6C).

### Validation of RNAi screen results using viable homozygous mutants

Lastly, we verified the RNAi screen results using available mutant strains capable of producing embryos as homozygotes. We tested the requirement of *dhhc-9*, *sem-5*, *ZK637.2*, and *pgp-3* in asymmetric ring closure in zygotes and found that the *dhhc-9(gk985)* deletion mutant and *sem-5(cy15)* mutant (a premature stop codon at amino acid 192 a.a., resulting in loss of the C-terminal 192–228 a.a.) exhibited a reduced rate of asymmetric furrowing (Figure 7A). We also tested the requirement of *sem-5*, *pyr-1*, *bmk-1*, *dod-20*, and *rgl-1* in the rotation of the AB cell at the two-cell stage and found that the *sem-5(cy15)* mutant exhibited a reduced degree of cellular rotation (Figure 7B). Finally, we tested the requirement of *mak-1*, *ufd-3*, and *nab-1* in the posterior shift of the cleavage furrow during EMS cell division and found that the *mak-1(ok2987)* deletion mutant exhibited an extensive posterior furrow shift (Figure 7C).

**Figure 7.**
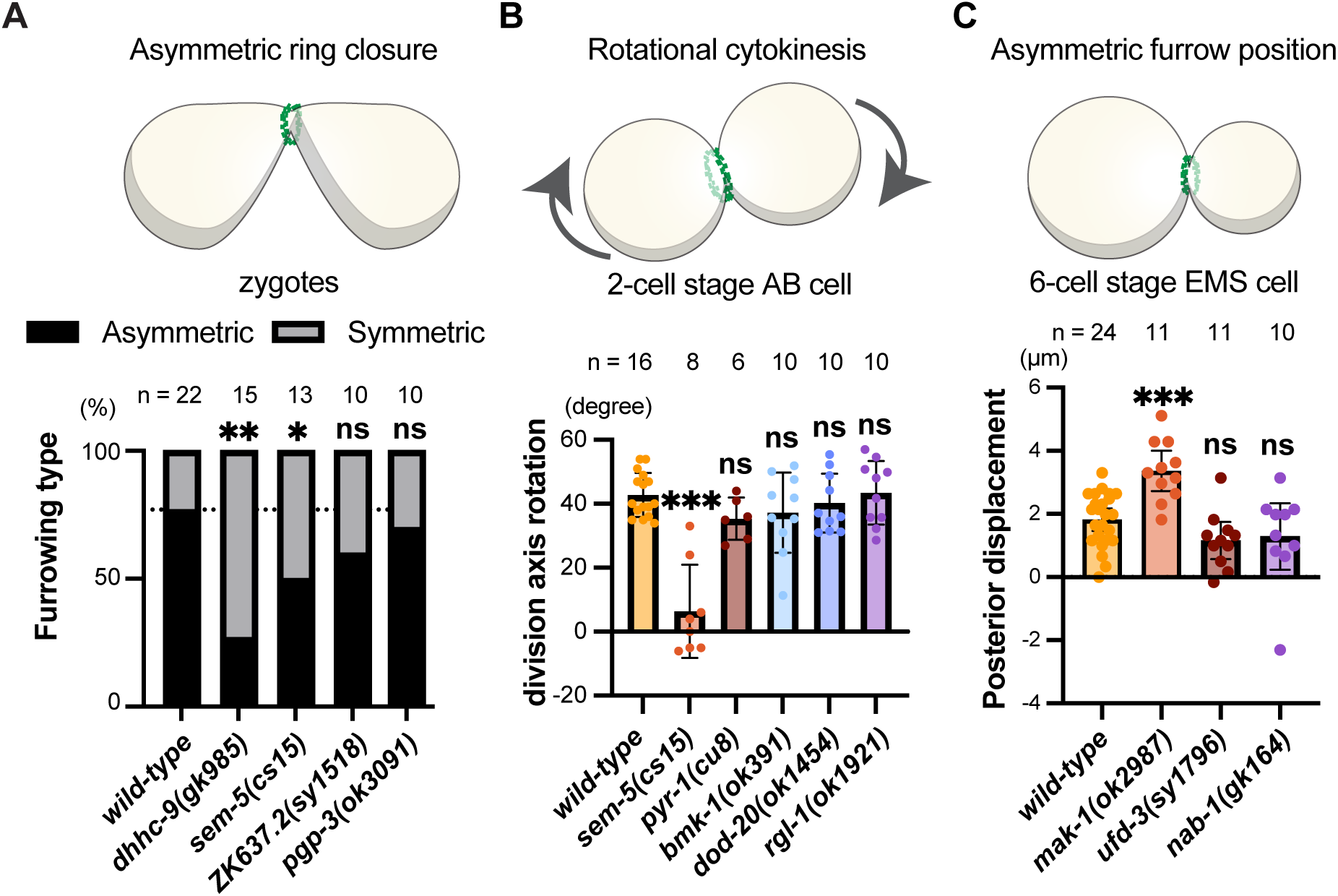
RNAi screen results were partially confirmed in mutant analyses. (**A**-**C**) Cytokinesis phenotypes in viable homozygous mutants. Phenotypes were quantified for asymmetric furrowing in zygotes (A), the angle of cellular rotation in the two-cell stage AB cell (B), and the posterior shift of the cleavage furrow in the six-cell stage EMS cell (C), as described in Figure 1. The same wild-type dataset shown in Figure 1 was used for comparison. Error bars represent 95% confidence intervals. Fisher’s exact test was used in panel A, and Brown-Frosythe and Welch ANOVA tests were used in panels B and C. P-values: “***”, “**”, “*”and “ns” indicate p < 0.001, p < 0.01, p < 0.05, and p > 0.05, respectively.

## Discussion

In this study, we found that the gain-of-function actin mutant *act-2(or295)* exhibits defects in the spatial control of cytokinesis. By leveraging this genetic background, we conducted a high-throughput RNAi screen and identified 689 suppressors of *act-2(or295)* embryonic lethality. In a secondary, live-imaging-based screen in a wild-type background, we identified 35 genes involved in asymmetric contractile ring closure in zygotes, 16 genes involved in rotational cytokinesis in the 2-cell stage AB cell, and 21 genes involved in asymmetric contractile ring positioning in the 6-cell stage EMS cell. Furthermore, we conducted another secondary RNAi screen in *act-2(or295)* background to search for genes enabling successful cytokinesis under mechanical stress. We identified four genes that exhibit cytokinesis defects in *act-2(or295)*.

Although the position and function of the contractile ring are spatially controlled in tissues, the mechanisms underlying this spatial control remain unclear. Through our RNAi screen, we identified potential regulators for each of the three modes of spatial cytokinesis control. Among the 69 candidate genes, only three genes were common to at least two modes of spatial control. Some of the identified genes are known to be involved in cytokinesis. For example, we identified *dhhc-9*/ZDHHC16, which encodes S-palmitoyltransferase, as a regulator of asymmetric contractile ring closure (Figure 5A). A recent study in mammalian cultured cells shows that ZDHHC5 regulates the palmitoylation of protocadherin 7 (PCDH7) during mitosis, thereby targeting its cleavage furrow localization and activating RhoA (Ozkan *et al*. 2023). Disruption of PCDH7 increases cytokinesis failure, but its role in asymmetric contractile ring closure remains unexplored. We also identified *sem-5*/Grb2 as a regulator of rotational cytokinesis (Figure 5B). Grb2 is a protein containing Src homology domain 2 (SH2) and SH3 domains, acting as an adaptor in various cell signaling pathways, such as FGF and EGF signaling (Giubellino *et al*. 2008). Grb2 interacts with Wiskott-Aldrich Syndrome protein (WASp) and with cortactin, both of which regulate the branched actin nucleator Arp2/3 complex (She *et al*. 1997; Martinez-Quiles *et al*. 2004). Interestingly, our candidates for rotational cytokinesis regulators also include the N-WASp homolog *wsp-1*. The list of candidate genes generated in this study should inspire novel hypotheses by integrating this existing molecular knowledge.

The contractile ring is a molecular machine known for its remarkable resilience, yet the mechanisms underlying this resilience of cytokinesis are not yet fully understood. Our secondary screen in *act-2(or295)* mutants identified four genes that contribute to cytokinesis completion in this background. Genes, such as *tba-1*, *spd-2*, and *evl-20* are involved in microtubule assembly. The mitotic spindle plays a critical role in activating RhoA through the central spindle and astral microtubules (Pollard and O’Shaughnessy 2019). A previous small-scale RNAi screen in *C. elegans* also identified microtubule regulators as genes required for the mechanical resilience of cytokinesis (Singh *et al*. 2019). That experimental setup was more specific, subjecting embryos to uniaxial compression using small-diameter beads and a coverslip. Thus, our identification of these microtubule regulators confirms their requirement in the mechanical resilience of cytokinesis. We also identified the Lipin homolog *lpin-1* as a regulator of mechanical resilience of cytokinesis. The specific role of Lipin in contractile ring formation and function has yet to be determined and will be an interesting direction for future research.

It is worth noting that this study has four main limitations. First, the primary RNAi screen, which analyzed 4,218 early embryonic genes, was performed only once. Consequently, some pseudonegatives may have gone undetected due to technical errors. Second, both secondary RNAi screens used feeding RNAi with relatively small sample sizes, limiting our ability to fully elucidate the roles of candidate genes in cytokinesis. Third, although we verified the RNAi screen results using mutants, the analysis was limited to homozygous viable strains, as several available mutants were larval lethal or sterile and therefore did not produce embryos. Finally, *act-2(or295)* enhancers likely include other interesting candidates, but we did not analyze them in order to streamline the study.

Overall, this study identified a number of potentially novel regulators of cytokinesis, with a particular focus on the relatively underexplored areas of cytokinesis research, mechanical resilience and spatial control. The identified list of genes will facilitate the generation of future hypotheses aimed at understanding the multicellular-specific regulation of cytokinesis during animal development.

## Supporting information

List of act-2 suppressors

## Acknowledgements

We thank Caenorhabditis Genetics Center (funded by the NIH Office of Research Infrastructure Programs; P40 OD010440) for sharing worm strains and the Sugioka lab members for general discussions. We also thank Life Sciences Institute Imaging Core Facility, Michele Roberge, Aruna Balgi for sharing high-content screening microscopy, Don Moerman and Mark Edgley for sharing liquid handling machine. This work was supported by the Canadian Institutes of Health Research (Project Grant; PJT-169145), Government of Canada’s New Frontiers in Research Fund (NFRFE-2019-00310), and the Michael Smith Health Research BC (Scholar Award; SCH-2020-0406) to K.S.

## Author contributions

**C.G.**: Formal analysis, Investigation, Data curation, Writing-Original Draft, Validation, Visualization. **Y.R.X.**: Formal analysis, Investigation, Data Curation, Project administration. **A. H.**: Formal analysis, Investigation, Methodology, Data Curation, Project administration. **T.L.**: Investigation, Formal analysis. **C.R.H.**: Investigation. **V. J.**: Data Curation. **M.J.K.** Investigation. **K.D.** Investigation. **K.S.**: Conceptualization, Formal analysis, Investigation, Methodology, Validation, Writing-Original Draft, Writing-Review & Editing, Supervision, Funding acquisition.

## Declaration of interests

We declare no competing interests.

## Materials and Methods

### Maintenance of worm strains

All *C. elegans* strains were cultured and maintained using standard methods (Brenner 1974). The transgene, *oxTi872*[*vha-6p*::GFP::*tbb-2* 3’UTR + Cbr-*unc-119*(+)], was used to monitor intestinal GFP expression. Additionally, *act-2(or295)* mutants were used as our sensitized genetic background. Both the *act-2(or295)* mutants and wild-type strains carrying *oxTi872* were maintained on NGM agar plates at 15°C. During the liquid culture RNAi screen, these strains were incubated at 20°C, the semi-permissive temperature for *act-2(or295)* mutant strain. For both secondary screens, these strains were incubated at 25°C, the restrictive temperature for *act-2(or295)*.

For confocal microscopy imaging in Figure 2, strains carrying *nmy-2(cp13)*[*nmy-2*::GFP + LoxP] (knock-in reporter, (Dickinson et al. 2013)), with or without *act-2*(or295) mutation, were used. These strains were maintained at 20°C and imaged at room temperature (∼22.5°C).

The following homozygous viable mutant strains were used to validate the RNAi screen results: *dhhc-9*(gk985), *sem-5*(cy15), *ZK637.2*(sy1518), *pgp-3*(ok3091), *pyr-1*(cu8), *bmk-1*(ok391), *dod-20*(ok1454), *rgl-1*(ok1921), *mak-1*(ok2987), *ufd-3*(sy1796), and *nab-1*(gk164).

### RNAi

Feeding RNAi was performed using the Ahringer RNAi library (Fraser *et al*. 2000; Ahringer 2006). The expression of dsRNA in the *Escherichia coli* strain HT115 was induced by overnight incubation at 37°C in Nematode Growth Medium (NGM) containing 10 µg/mL Ampicillin and 1 mM isopropyl ß-D-1-thiogalactopyranoside (IPTG).

For the high-throughput RNAi screen, gravid adults were bleached with hypochlorite solution, and the resulting purified embryos were incubated in M9 buffer overnight to obtain developmentally arrested L1 stage worms. These worms were suspended in liquid NGM and approximately 4-5 L1 worms were transferred into each well of 96-well plates containing various dsRNA-expressing bacteria. After 5 days of incubation at 20°C, the 96-well plates were imaged using a high-content screening system (Cellomics Arrayscan; Thermo Fisher).

For the two secondary RNAi screens, dsRNA-expressing bacteria were cultured on solid NGM agar plates. Developmentally arrested L1 worms were transferred onto these plates and incubated for 3 days at 25°C. For genes required for fertility, we used L4 stage worms were used and incubated overnight before live imaging. Wild-type worms and *act-2(or295)* mutants were used for the spatial cytokinesis regulator screen and the cytokinesis resilience regulator screen, respectively. A bacterial strain containing the empty vector L4440 served as a negative control in both secondary screens.

In Figure 2, partial knockdown of ECT-2 was achieved by incubating *act-2*(or295) worms on IPTG-containing NGM agar plates at 20°C for 6 hours prior to imaging.

### Live-imaging

To obtain embryos for live imaging by brightfield microscopy, we dissected gravid adults on a coverslip in a 5 μl droplet of M9 buffer. We then placed the coverslip gently onto a 2% agarose pad mounted on a glass slide and sealed the edges with petroleum jelly (Vaseline) to prevent dehydration.

For characterization of the strains used in the high-throughput RNAi screen and secondary screens, we performed live imaging at room temperature (∼22.5°C) using brightfield compound microscopes (B290-TB; OPTIKA), equipped with a 100X oil objective lens (1.2NA) and controlled by OPTIKA Vision Lite software. For the secondary screen of spatial cytokinesis regulators, images were taken every 2 seconds for 50 minutes, beginning at the 1-cell stage and continuing until after EMS cell division. For the secondary screen of cytokinesis resilience regulators, images were taken every 2 seconds for 10 minutes, beginning at the 1-cell stage and ending at the 2-cell stage. In most cases, filming began during late pronuclear migration stage but before nuclear envelope breakdown (NEBD).

For confocal microscopy imaging of wild-tpe and act-2(or295), dissected embryos were mounted on 2% agarose and imaged using an Olympus IX83 microscope (Olympus/Evident), equipped with a spinning-disk confocal unit CSU-W1 (Yokogawa), a scientific CMOS camera Prime95B (Photometrics), a piezoelectric stage NANO-Z (Mad City Labs), a silicon immersion objective UPLSAPO60XS2 (NA1.3, 60X; Olympus/Evident), all controlled by Cellsense Dimension software (Olympus/Evident). Silicone immersion oil (Z81114; refractive index: 1.406 at 23 °C; Olympus/Evident) was used as the immersion medium. Imaging of myosin II::GFP was performed with 488 nm diode-pumped lasers, with settings of 150 ms exposure time, 1 µm Z-step size, 28 slices per frame, and a 5.57 second frame interval.

### Image analysis for the high-throughput RNAi screen

To quantify the number of hatched worms, we processed images obtained from the high-content screening system using Fiji (Schindelin *et al*. 2012). We applied a Laplacian filter to extract edge information while removing background signals. Next, we used Otsu’s threshold to identify relevant signals and then applied morphological filters to select rod-like worm bodies while excluding circular objects, such as embryos and debris. The resulting segmentation data was used to count the number of hatched F1 and F2 worms. We then standardized the worm counts for each 96-well plate by calculating Z-scores, using the mean and standard deviation of worm counts within each plate to minimize batch effects. We computed three different Z-scores for each candidate gene: one in the control background, one in the *act-2(or295)* mutant background, and the difference between the two. Genes with Z-scores greater than 1.5 in the *act-2(or295)* mutant background were considered actin suppressors. Additionally, genes with a Z-score difference greater than 1.5 between *act-2* and control strains were also designated as actin suppressors.

### Selection of candidate genes for secondary screens

For the secondary screen of spatial regulators of cytokinesis, all actin suppressor genes were assessed using plate-based RNAi in a wild-type strain (EG8829). At least three embryos were imaged for each gene, and phenotypes were manually annotated in Fiji using the collected live-imaging movies.

For the secondary screen of regulators of cytokinesis resilience, we first selected actin suppressor genes using two different criteria. Initially, we chose genes based on defects in asymmetric ring closure in 1-cell stage zygotes, as identified in the spatial cytokinesis regulator screen. However, we found that most of these genes were not required for cytokinesis in the *act-2* mutant background. Therefore, we selected genes whose RNAi knockdown cased 100% embryonic lethality in *act-2* mutants at 25°C for genes on chromosomes III-X.

### Bioinformatics

Scatter plots for the high-throughput screening data were generated using Matplotlib. To analyze and visualize gene networks among the identified candidates, we used GeneMANIA in the Cytoscape software (Shannon *et al*. 2003).

### Statistics

Statistical analyses were performed using Prism9 (GraphPad Software) and custom Python scripts. For Figure 1E, we performed Fisher’s exact test on raw data (rather than percentages). For Figure 1F-G, we used Welch’s t-test. For Figure 6B, we used Fisher’s exact test on raw data. The resulting P-values were corrected for multiple comparisons using a custom Python script (Holm-Sidak method). The symbols such as ****, ***, **, *, and ns indicate p < 0.0001, p < 0.001, p < 0.01, p < 0.05, and p > 0.05, respectively.

## Notes

### Competing Interest Statement

The authors have declared no competing interest.

### Summary of Updates

We have added confocal microscopy data in Figure 2. Also, Figure 7 using mutants were added.

